# Physiological Oscillations of Lateral Ventricular Volume Revealed by Fast fMRI: Effects of Aging and Vigilance

**DOI:** 10.64898/2026.10.24.761584

**Authors:** Leiyu Wang, Adam M. Wright, Yi Zhao, Yunjie Tong, Jason W. Allen, Qiuting Wen

## Abstract

**Background:** Cyclic ventricular deformation may contribute to CSF displacement. However, the physiological mechanisms underlying ventricular motion remain poorly understood. This study characterized physiological oscillations of lateral ventricular volume (LVV) and examined their associations with aging and vigilance.

**Methods:** We analyzed two fast resting-state fMRI scans from 99 adults aged 35–82 years. The first scan was acquired during eyes-open wakefulness, followed by 30 min of eyes-closed rest and a second scan during eyes-closed drowsiness. LVV was estimated for each image volume using automated ventricular segmentation, yielding time-resolved volume measurements. Temporal variability was quantified using the standard deviation and coefficient of variation, and spectral power was quantified for low-frequency, respiratory, and cardiac oscillations. Age-related associations were assessed using multiple linear regression adjusted for sex. Awake–drowsy differences were assessed using paired t-tests. False discovery rate correction was applied for multiple comparisons in all analyses.

**Results:** After quality-control exclusions, 97 participants were analyzed. LVV exhibited distinct low-frequency, respiratory, and cardiac oscillatory components. Respiratory and cardiac oscillations accounted for 37.7% and 38.4% of total physiological spectral power, respectively, compared with 23.9% for low-frequency oscillations. Mean lateral ventricular volume increased with age (FDR-adjusted p = 0.010), while respiratory and cardiac power also showed positive associations with age, although neither survived FDR correction (both FDR-adjusted p = 0.067). The transition to eyes-closed drowsiness was associated with greater LVV fluctuation magnitude, reflected by increases in standard deviation (FDR-adjusted p < 0.001) and coefficient of variation (FDR-adjusted p = 0.028) and accompanied by increased low-frequency (FDR-adjusted p < 0.001) and respiratory power (FDR-adjusted p = 0.015). Mean LVV and cardiac power did not change significantly between conditions.

**Conclusions:** LVV exhibits prominent respiratory and cardiac oscillations in addition to low-frequency fluctuations, suggesting that ventricular dynamics reflect multiple physiological influences rather than a single dominant physiological driver. Aging was primarily associated with ventricular enlargement, whereas reduced vigilance was accompanied by increased LVV fluctuation magnitude, primarily reflected in increased low-frequency and respiratory power, without a change in mean LVV. Time-resolved LVV provides a complementary approach for investigating the physiological regulation of ventricular dynamics in health and disease.

## 1. Introduction

Cerebrospinal fluid (CSF) circulates through the ventricular and subarachnoid spaces and exchanges with fluid in the brain parenchyma, supporting brain homeostasis and waste removal [1–5]. Cyclic deformation of the ventricular walls may contribute to CSF displacement and circulation. The lateral ventricles are particularly relevant because they are the largest ventricular compartments and contain substantial CSF-producing choroid plexus, making their rhythmic deformation potentially important for CSF displacement and redistribution. However, the physiological forces underlying lateral ventricular wall motion in humans remain poorly understood. Cyclic vascular and pressure fluctuations may contribute to ventricular deformation and lateral ventricular volume (LVV) fluctuations [1,6]. Low-frequency oscillations (LFOs), respiration, and cardiac pulsation have each been implicated in human CSF dynamics [7–10], but their relative contributions to LVV fluctuations remain unclear.

Previous human studies have identified different dominant physiological drivers of ventricular CSF oscillations. Earlier cardiac-gated and CINE MRI studies also demonstrated dynamic changes in lateral ventricular volume across the cardiac cycle in healthy individuals [11,12]. Some have emphasized LFOs [10,13–15], whereas others have emphasized cardiac [10,13] or respiratory effects [7,8,10]. These differences may reflect variations in imaging contrast and the CSF compartments examined, with some studies quantifying CSF flow in the cerebral aqueduct and others measuring inflow-sensitive CSF signal fluctuations in the fourth ventricle. Because ventricular CSF motion is mechanically coupled to deformation of the ventricular system [16], time-resolved LVV may provide a more direct assessment of the lateral ventricles’ mechanical response to physiological oscillations.

Recent advances in fast fMRI acquisition and automated brain segmentation enable LVV to be estimated for each acquired image volume [17–19], allowing time-resolved quantification across low-frequency, respiratory, and cardiac frequency bands. This approach also provides an opportunity to examine how LVV dynamics vary with aging and transient changes in vigilance. Aging is associated with alterations in ventricular morphology and CSF physiology [20,21], whereas reduced vigilance is accompanied by shifts in neural and autonomic activity that may alter forces acting on the ventricular system [10,15,16,22]. However, the effects of aging and vigilance on LVV dynamics remain poorly understood.

In the present study, we combined fast resting-state fMRI with an automated LVV segmentation approach to characterize LVV dynamics in healthy adults. We aimed to (1) quantify LVV fluctuations in both the time and frequency domains and assess the relative contributions of low-frequency, respiratory, and cardiac oscillations to LVV dynamics, (2) determine whether LVV metrics differ with age, and (3) examine state-related differences in LVV dynamics between eyes-open awake and eyes-closed drowsy conditions.

## 2. Methods

### 2.1 Participants and study design

Data from a total of 99 healthy participants (69 female, 30 male) aged 35–82 years (53.31 ± 12.85 years) were initially included in this study. Each participant completed two resting-state fMRI scans during the same imaging session. The first scan was acquired with participants instructed to keep their eyes open and remain still and was designated the awake condition. Participants then underwent a 30-minute uninterrupted eyes-closed rest period intended to promote drowsiness. After that, a second resting-state fMRI scan was acquired while participants kept their eyes closed and was designated the drowsy condition. Following imaging, participants retrospectively rated their subjective level of drowsiness during each conditions using the 8-point Stanford Sleepiness Scale [23,24], with ranging from 1 (full alertness) to 8 (asleep). The study was approved by the Indiana University Institutional Review Board and was conducted in accordance with the Declaration of Helsinki. Written informed consent was obtained from all participants before participation.

### 2.2 MRI acquisition

All imaging data were acquired on a 3T Siemens Prisma MRI scanner equipped with a 64-channel head-neck coil. The imaging protocol included a high-resolution structural T1-weighted MPRAGE scan (TR/TE = 2300/2.98 ms, flip angle = 9°, voxel size = 1.0 × 1.0 × 1.0 mm^3^), followed by two resting-state functional MRI (fMRI) scans acquired using a fast echo-planar imaging sequence (TR/TE = 363/30 ms, flip angle = 35°, voxel size = 2.5 × 2.5 × 2.5 mm^3^, 500 volumes, scan duration = 3 min 2 s per scan). Physiological signals, including respiratory belt measurements and finger pulse plethysmography, were recorded simultaneously during both fMRI scans using the Siemens built-in physiological monitoring system. These recordings were used to determine subject-specific respiratory and cardiac frequencies for subsequent analyses of physiological contributions to LVV dynamics.

### 2.3 LVV extraction and quality control

The fMRI data were first denoised using overcomplete local principal component analysis (LPCA). SynthSeg was then used to generate ventricular masks for each denoised fMRI volume and estimate LVV at each time point, yielding a time-resolved LVV time series [18,19]. The first 10 fMRI volumes were discarded to allow signal stabilization. The resulting LVV time series were demeaned and linearly detrended before subsequent time- and frequency-domain analyses. Segmentation quality was assessed using the General CSF quality control (QC) score generated by SynthSeg, which ranges from 0 to 1, with higher values indicating greater confidence in segmentation quality [19,25]. A QC score of 0.65 or higher was considered indicative of acceptable segmentation quality. Segmentations with QC scores below 0.65 underwent visual inspection to assess segmentation accuracy [19,25]. Participants with mean QC scores below 0.65 were excluded from subsequent analyses.

### 2.4 Quantification of LVV dynamics

LVV dynamics were quantified using both time- and frequency-domain metrics. Time-domain metrics included the mean LVV, the standard deviation (SD) of the LVV time series, and the coefficient of variation (CV), calculated as the SD divided by the mean LVV and expressed as a percentage.

Frequency-domain metrics were quantified using power spectral density (PSD) analysis. PSD was estimated from each LVV time series using Welch’s method (MATLAB pwelch function; Hanning window = 128 points, 50% overlap, nFFT = 512) [26]. Three physiological frequency components were evaluated: low-frequency oscillations (LFOs; 0.01–0.10 Hz), respiratory oscillations (respiratory peak frequency ± 0.10 Hz), and cardiac oscillations (cardiac peak frequency ± 0.10 Hz). Respiratory and cardiac peak frequencies were identified from the simultaneously recorded physiological signals for each participant. Absolute power was calculated by integrating the PSD within each frequency range. Relative power for each physiological component was calculated as its absolute power divided by the sum of the absolute powers across the LFO, respiratory, and cardiac bands, multiplied by 100% [27].

### 2.5 Statistical analysis

To investigate age-related effects on LVV dynamics, multiple linear regression models were constructed for each LVV metric, including mean LVV, SD, CV, LFO power, respiratory power, and cardiac power, using the data from the eye-open awake scan. Age and sex were included as independent variables in all regression models. Regression coefficients (β), 95% confidence intervals (CI), p-values, and model R^2^values were reported. To investigate within-subject differences between the eyes-open awake and eyes-closed drowsy conditions, paired t-tests were used to compare each LVV metric between conditions. Mean differences, Cohen’s d effect sizes, 95% confidence intervals, and p-values were calculated for all paired comparisons. For each analysis family, p-values were corrected for multiple comparisons across the six LVV metrics (mean LVV, SD, CV, LFO power, respiratory power, and cardiac power) using the Benjamini–Hochberg false discovery rate (FDR) procedure [30]. Statistical significance was defined as a two-sided FDR-adjusted p < 0.05. All statistical analyses were performed in R (version 4.6.1; R Foundation for Statistical Computing, Vienna, Austria).

## 3. Results

### 3.1 Temporal and spectral characteristics of LVV fluctuations during eyes-open rest

Of the 99 participants initially included, two were excluded because their mean QC scores were below the prespecified threshold, yielding a final sample of 97 participants (67 females and 30 males; mean age, 53.42 ± 12.55 years). During eyes-open rest, mean LVV during eyes-open rest was 14.95 ± 7.20 cm^3^, and LVV SD was 1.67 ± 1.06 cm^3^. The median LVV CV was 8.17% (IQR, 3.58%–16.61).

A representative lateral ventricle segmentation, the corresponding LVV time series, and its power spectrum are shown in Figure 1AB. The LVV power spectrum demonstrated three distinct oscillatory components corresponding to LFO, respiration, and cardiac pulsation (Figure 1B). To compare the relative contributions of these physiological components to overall LVV fluctuations, power in each frequency band was expressed as a percentage of the summed power across the three bands. At the group level, respiratory and cardiac oscillations were the two largest contributors to LVV fluctuations, accounting for 37.71 ± 13.76% and 38.43 ± 15.54% of the summed power across the three physiological components, whereas LFOs accounted for 23.87 ± 14.67% (Figure 1C). The relative contributions of the three physiological components differed significantly (Friedman test: χ^2^(2) = 42.687, *p* < 0.001, Kendall’s *W* = 0.216). Post hoc pairwise comparisons showed greater respiratory and cardiac oscillations than LFOs (both FDR-adjusted *p* < 0.001), whereas respiratory and cardiac contributions did not differ significantly (FDR-adjusted *p* = 0.710).

**Figure 1.**
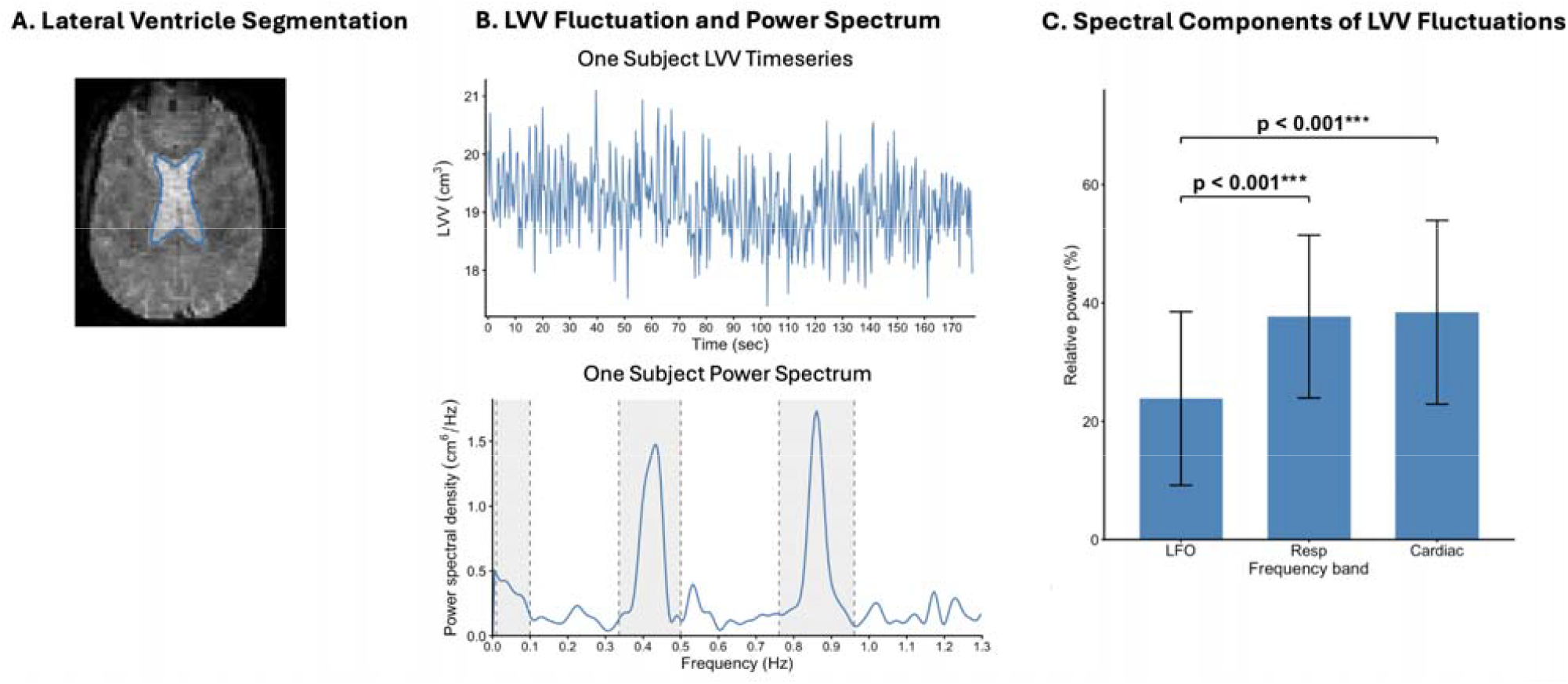
Physiological components of LVV fluctuations during eyes-open rest. (A) Representative lateral ventricle segmentation from a single fMRI volume using SynthSeg. (B) Representative LVV time series and corresponding power spectrum from one participant. Gray shaded regions indicate the frequency bands used for spectral quantification. (C) Group-level relative power of the three physiological components, showing greater respiratory and cardiac contributions than LFOs. Error bars indicate SD.

### 3.2 Age- and sex-related changes in LVV dynamics

Absolute power was used to assess age-related changes in the magnitude of each physiological oscillation independently of changes in the other frequency bands. Older age was associated with greater mean LVV (β = 0.227 cm^3^/year, 95% CI: 0.088–0.366, R^2^= 0.214, *p* = 0.002, FDR-adjusted *p* = 0.010, Figure 2 and Table 1). Older age was also nominally associated with higher respiratory power (β = 0.012 cm /year, 95% CI: 0.001–0.024, R^2^= 0.144, *p* = 0.033, FDR-adjusted *p* = 0.067) and higher cardiac power (β = 0.014 cm /year, 95% CI: 0.002–0.027, R^2^= 0.131, *p* = 0.025, FDR-adjusted *p* = 0.067), although neither association survived FDR correction. No significant associations were observed between age and LVV SD, CV, or LFO power (all FDR-adjusted *p* > 0.05).

**Table 1.**
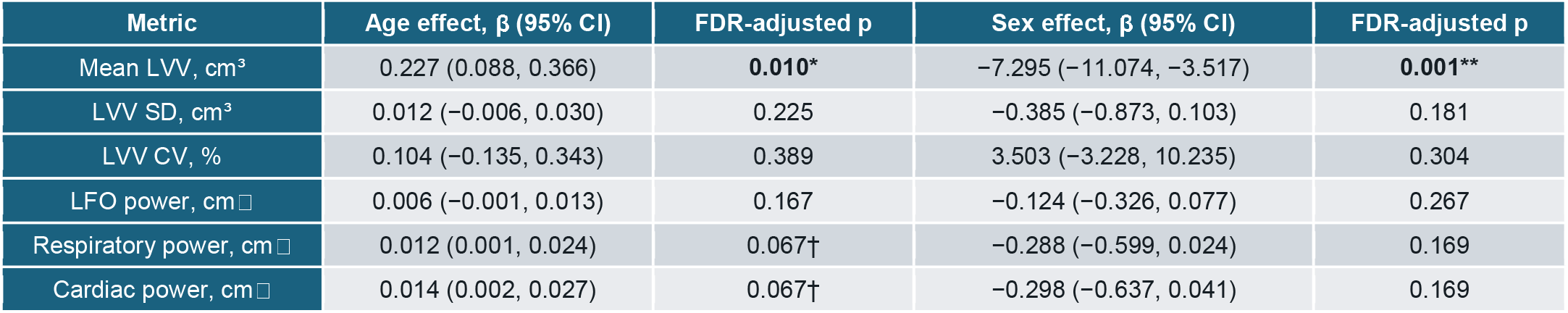
Associations of age and sex with LVV metrics. Values are regression coefficients (β) with 95% confidence intervals (CI). Sex effects represent females relative to males, with males as the reference group. p values were adjusted for multiple comparisons using the Benjamini–Hochberg false discovery rate (FDR) procedure. † FDR-adjusted p < 0.10; * FDR-adjusted p < 0.05; ** FDR-adjusted p < 0.01. Abbreviations: LVV, lateral ventricular volume; SD, standard deviation; CV, coefficient of variation; LFO, low-frequency oscillation; CI, confidence interval; FDR, false discovery rate.

| Metric | Age effect, $\beta$ (95% CI) | FDR-adjusted p | Sex effect, $\beta$ (95% CI) | FDR-adjusted p |
| --- | --- | --- | --- | --- |
| Mean LVV, cm<sup>3</sup> | 0.227 (0.088, 0.366) | **0.010\*** | -7.295 (-11.074, -3.517) | **0.001\*\*** |
| LVV SD, cm<sup>3</sup> | 0.012 (-0.006, 0.030) | 0.225 | -0.385 (-0.873, 0.103) | 0.181 |
| LVV CV, % | 0.104 (-0.135, 0.343) | 0.389 | 3.503 (-3.228, 10.235) | 0.304 |
| LFO power, cm<sup>2</sup> | 0.006 (-0.001, 0.013) | 0.167 | -0.124 (-0.326, 0.077) | 0.267 |
| Respiratory power, cm<sup>2</sup> | 0.012 (0.001, 0.024) | 0.067† | -0.288 (-0.599, 0.024) | 0.169 |
| Cardiac power, cm<sup>2</sup> | 0.014 (0.002, 0.027) | 0.067† | -0.298 (-0.637, 0.041) | 0.169 |

**Figure 2.**
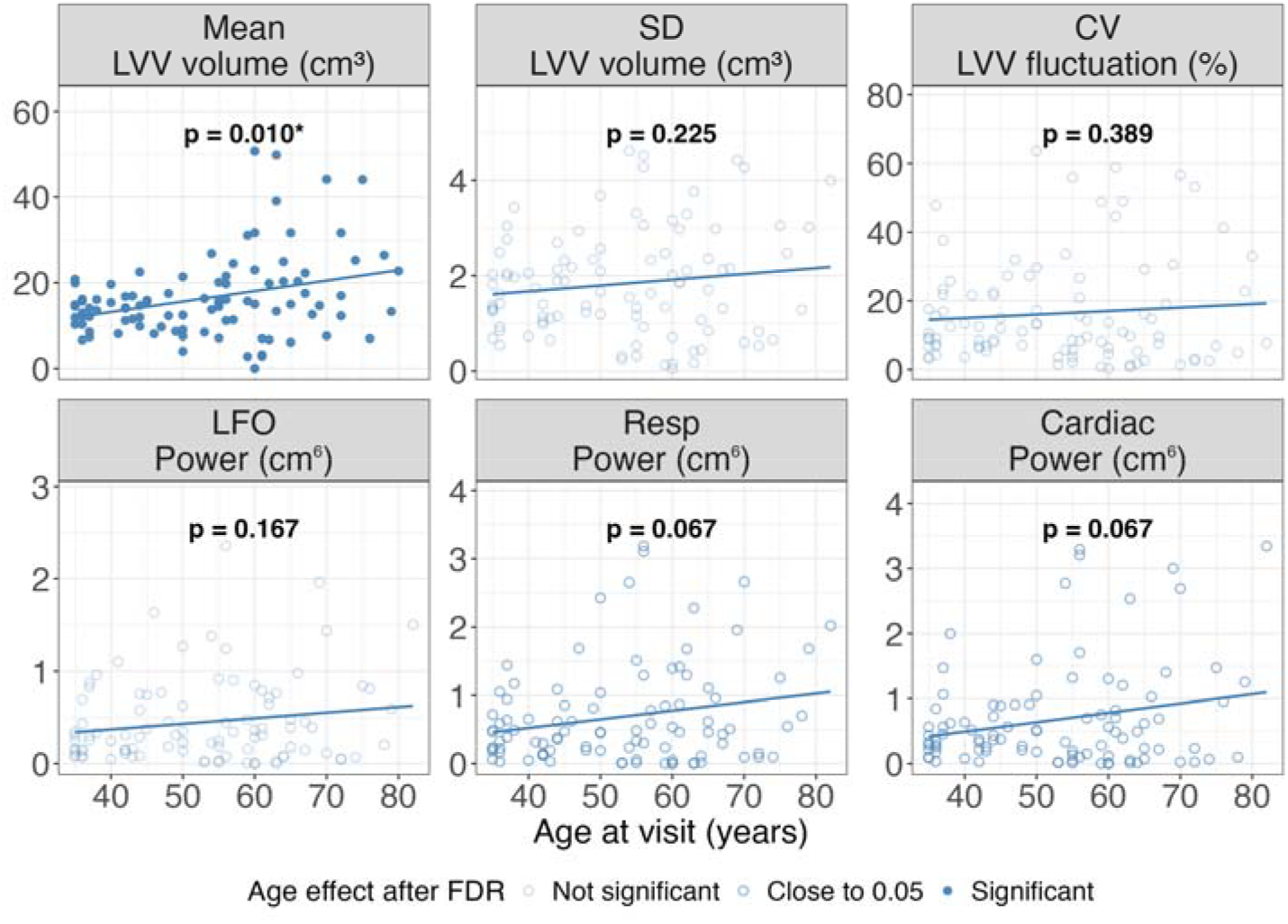
Associations between age and LVV metrics. Age was positively associated with mean LVV and nominally associated with respiratory and cardiac power, although the latter associations did not survive FDR correction. Solid lines represent linear regression fits. All p-values are FDR-corrected. * FDR-adjusted p<0.05; . FDR-adjusted p<0.1. Abbreviations: LVV, lateral ventricular volume; SD, standard deviation; CV, coefficient of variation; LFO, low-frequency oscillation; Resp, respiratory; FDR, false discovery rate.

After adjusting for age, females had lower mean LVV than males (β = −7.295 cm^3^, 95% CI: −11.074 to −3.517, FDR-adjusted *p* = 0.001; Table 1). No significant sex effects were observed for LVV SD, CV, LFO power, respiratory power, or cardiac power (all FDR-adjusted *p* > 0.05).

### 3.3 Changes in LVV dynamics from eyes-open wakefulness to eyes-closed drowsiness

Subjective sleepiness scores and physiological measures supported a transition toward reduced vigilance between the awake and drowsy conditions. SSS significantly increased from 4.52 ± 1.65 during the awake condition to 5.73 ± 1.97 during the drowsy condition (Cohen’s *d* = 0.52, FDR-adjusted *p* < 0.001), accompanied by a modest decrease in heart rate from 64.98 ± 9.11 to 64.02 ± 9.07 bpm (Cohen’s *d* = −0.26, FDR-adjusted *p* = 0.047) and an increase in heart rate variability, quantified using the standard deviation of normal-to-normal intervals (SDNN), from 39.93 ± 18.00 to 52.28 ± 23.71 ms (Cohen’s *d* = 0.57, FDR-adjusted *p* < 0.001). Respiratory rate did not differ significantly between conditions (14.28 ± 3.46 vs. 13.81 ± 3.30 breaths/min; Cohen’s *d* = −0.16, FDR-adjusted *p* = 0.180).

In parallel with these physiological changes, LVV dynamics also changed during drowsiness. Mean LVV did not differ between the awake and drowsy conditions (14.95 ± 7.20 vs. 14.63 ± 6.55 cm^3^, Cohen’s *d* = −0.12, FDR-adjusted *p* = 0.252). In contrast, LVV SD increased from 1.67 ± 1.06 to 1.93 ± 1.19 cm^3^ (Cohen’s *d* = 0.42, FDR-adjusted *p* < 0.001), and CV increased from 11.34 ± 9.55% to 12.88 ± 9.73% (Cohen’s *d* = 0.26, FDR-adjusted *p* = 0.028). Frequency-domain analysis showed increased LFO power (0.260 ± 0.277 vs. 0.408 ± 0.374 cm, Cohen’s *d* = 0.49, FDR-adjusted *p* < 0.001) and respiratory power (0.477 ± 0.470 vs. 0.620 ± 0.641 cm, Cohen’s *d* = 0.29, FDR-adjusted *p* = 0.015), whereas cardiac power did not differ significantly between the two conditions (0.438 ± 0.432 vs. 0.488 ± 0.443 cm, Cohen’s *d* = 0.18, FDR-adjusted *p* = 0.118).

**Figure 3.**
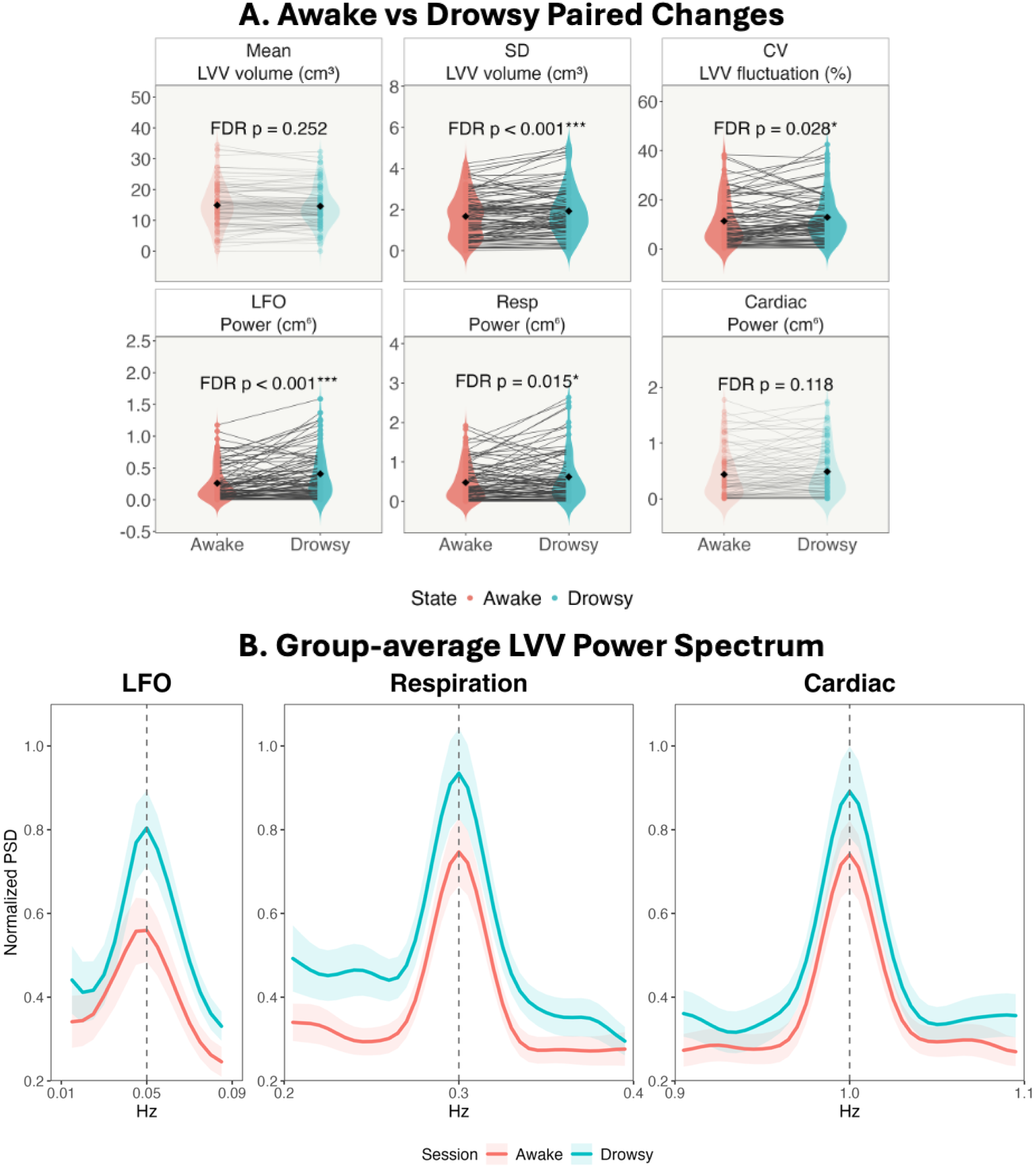
Changes in LVV dynamics between the awake and drowsy states. (A) Paired comparisons of LVV metrics between the awake and drowsy scans. Violin plots show the distribution of each metric, black diamonds indicate group means, and gray lines connect paired observations from individual participants. (B) Group-average normalized LVV power spectra for the awake and drowsy conditions. For group averaging, subject-specific respiratory and cardiac spectra were frequency-aligned to common reference peaks at 0.3 and 1.0 Hz, respectively, to minimize spectral blurring from interindividual frequency variability. Drowsiness was associated with increased LVV fluctuation amplitude, particularly in the LFO and respiratory frequency ranges. Abbreviations: LVV, lateral ventricular volume; SD, standard deviation; CV, coefficient of variation; LFO, low-frequency oscillation; PSD, power spectral density; FDR, false discovery rate.

## 4. Discussion

### 4.1 Summary of primary findings

In this study, we characterized LVV dynamics using fast fMRI and automated ventricular segmentation. LVV exhibited distinct low-frequency, respiratory, and cardiac oscillatory components, with respiratory and cardiac oscillations contributing the largest proportions of LVV power. With aging, mean LVV increased, while respiratory and cardiac power also tended to increase, although these associations did not survive correction for multiple comparisons. In contrast, reduced vigilance was accompanied by increased LVV fluctuation amplitude, particularly within the low-frequency and respiratory frequency ranges, without a significant effect on mean LVV. Together, these findings suggest that LVV dynamics provide a sensitive measure of physiological influences on ventricular motion, revealing distinct contributions of respiratory and cardiac oscillations and differential modulation by aging and vigilance.

### 4.2 Respiratory and cardiac oscillations are important contributors to LVV dynamics

Previous studies have implicated LFOs as an important contributor to ventricular CSF dynamics, particularly gBOLD-CSF coupling fMRI studies in which CSF dynamics are inferred from inflow-related signal changes in the fourth ventricle or spinal canal [10,13–15,28]. Our finding of a distinct LFO component within LVV dynamics is consistent with these observations. However, our findings extend the current literature by showing that respiratory and cardiac oscillations account for 37.7% and 38.4% of LVV fluctuations compared with 23.9% for LFOs. This suggests that lateral ventricular pulsation is driven by multiple physiological processes rather than predominantly by low-frequency oscillations. Our findings also complement previous phase-contrast MRI studies, which showed that CSF flow through the cerebral aqueduct and spinal canal is primarily cardiac driven [7,8,29–33]. While we likewise observed a strong cardiac component, respiration contributed a comparable proportion of LVV power. To our knowledge, the contribution of respiration to lateral ventricular volume dynamics has not been well characterized previously. This finding highlights respiration as a comparably important contributor to lateral ventricular dynamics at rest and suggests that the physiological determinants of ventricular wall motion may differ from those of CSF flow measured in the cerebral aqueduct and spinal canal.

The LFO, respiratory, and cardiac components were separated spectrally for quantification but should not be interpreted as physiologically independent drivers of ventricular motion. Respiration interacts with cardiovascular regulation through modulation of heart rate, venous return, blood pressure, and autonomic reflexes [34], while respiratory changes in arterial CO□ modulate cerebral vascular tone at lower frequency [35]. Cardiorespiratory oscillations are therefore interconnected across multiple physiological time scales [10,22]. Within the constrained intracranial space, these coupled vascular and pressure fluctuations in cerebral blood volume may be transmitted to the ventricular and CSF systems through shared intracranial volume-pressure mechanisms [1,6]. Thus, the spectral components observed in LVV may represent different temporal expressions of an integrated cardiorespiratory-cerebrovascular system rather than isolated physiological processes.

### 4.3 Aging influences ventricular size and physiological oscillations

Age-related ventricular enlargement is a well-established finding and was confirmed in the present study [20,36–38]. In contrast, age-related changes in LVV dynamics were less pronounced. LVV standard deviation, coefficient of variation, and LFO power showed no significant associations with age, whereas respiratory and cardiac LVV power showed positive trends that did not survive correction for multiple comparisons. Together, these findings suggest that aging has a much stronger effect on ventricular size than on short-timescale ventricular dynamics, although respiratory and cardiac components may show modest age-related increases.

The positive age associations observed for respiratory and cardiac LVV power may reflect age-related changes in the transmission of physiological pulsations to the ventricular system. Vascular stiffening and reduced vascular compliance with aging could diminish the buffering of cardiac and respiratory pulsations, potentially increasing their transmission through intracranial vascular and pressure fluctuations. The direction of these trends is consistent with recent fMRI studies reporting age-related increases in respiratory- and cardiac-related physiological oscillations across BOLD signals from brain tissue and vascular compartments [39]. Together, these observations support the possibility that age-related changes in cerebral vascular pulsatility may be reflected in ventricular motion, although this interpretation remains tentative because the respiratory and cardiac associations did not survive correction for multiple comparisons.

### 4.4 Reduced vigilance enhances LVV fluctuations

Reduced vigilance was accompanied by greater LVV fluctuation amplitude without significantly changing mean LVV, indicating that vigilance primarily modulates ventricular dynamics rather than ventricular size. Both LVV standard deviation and coefficient of variation increased during drowsiness, and frequency-domain analysis showed that this increase was primarily reflected in greater LFO and respiratory power, whereas cardiac power remained unchanged. This pattern is consistent with previous studies reporting enhanced low-frequency neurofluid dynamics during reduced vigilance and sleep [10,15,16,28,31], as well as magnetic resonance encephalography (MREG) studies showing increased respiratory power with reduced vigilance [22,40,41]. Our results extend these observations by demonstrating that similar state-dependent cerebral physiological changes are expressed directly in lateral ventricular volume fluctuations.

The increase in LFO power may reflect broader changes in autonomic and vascular regulation that accompany the transition toward drowsiness. In the present study, heart rate decreased modestly while heart rate variability, measured by SDNN, increased substantially, consistent with known shifts in cardiovascular autonomic regulation during reduced vigilance and sleep onset [42,43]. Because autonomic cardiovascular regulation and low-frequency vascular oscillations are closely interconnected [10,42,43], the concurrent increase in heart rate variability and LFO power raises the possibility that enhanced low-frequency ventricular fluctuations reflect, at least in part, a change in autonomic–vascular activity during drowsiness. Respiratory rate did not change significantly between conditions despite a significant increase in LVV respiratory power. This suggests that the stronger respiratory component cannot be explained simply by a change in breathing frequency. Instead, reduced vigilance may alter the transmission of respiratory-related physiological fluctuations to the ventricular system. One possible mechanism is a state-dependent change in brain tissue and interstitial properties; studies have shown that sleep is associated with expansion of the interstitial space and reduced resistance to fluid movement [44,45], which could facilitate the transmission of physiological pulsations. Together, these findings suggest that drowsiness enhances ventricular pulsatility rather than altering ventricular size, and that this enhancement may reflect coordinated changes in low-frequency vascular, respiratory, and tissue mechanical processes.

### 4.5 Limitations

This study has several limitations. First, the spatial resolution of the fMRI acquisition (2.5 × 2.5 × 2.5 mm^3^) limits the precision of LVV estimation, particularly for detecting small changes in ventricular volume. As a reference, a one-voxel dilation of the ventricular mask increased ventricular volume by 34.80 ± 9.40% across participants, substantially greater than the observed LVV fluctuations of 11.34 ± 39.55%. The magnitude of the observed LVV fluctuations therefore likely reflects small, potentially sub-voxel-scale changes in the ventricular boundaries. Higher-resolution imaging may improve the accuracy of LVV measurements and characterization of subtle physiological fluctuations, although this would come at the expense of temporal resolution.

Second, reduced vigilance was inferred from subjective sleepiness score and concurrent physiological measurements rather than objectively confirmed using EEG-confirmed sleep staging. Future studies combining fast fMRI with simultaneous EEG could provide more precise characterization of vigilance-dependent changes in LVV dynamics.

### 4.6 Conclusions

Together, these findings indicate that LVV dynamics capture distinct physiological influences on ventricular motion and are sensitive to state-dependent changes. Time-resolved LVV complements approaches based on CSF flow and gBOLD dynamics and can be extracted automatically from fast resting-state fMRI without requiring a separate acquisition. This approach may provide a practical framework for investigating the physiological regulation of ventricular dynamics in health and disease.

## List of abbreviations

CSF: Cerebrospinal fluid
CV: Coefficient of variation
FDR: False discovery rate
fMRI: Functional magnetic resonance imaging
LFO: Low-frequency oscillation
LVV: Lateral ventricular volume
MPRAGE: Magnetization-prepared rapid gradient echo
MRI: Magnetic resonance imaging
PSD: Power spectral density
QC: Quality control
SD: Standard deviation
SDNN: Standard deviation of normal-to-normal intervals
SSS: Stanford Sleepiness Scale
TR: Repetition time
TE: Echo time

## Declarations

### Ethics approval and consent to participate

The study was approved by the Indiana University Institutional Review Board ((Protocol No. 19788) and was conducted in accordance with the Declaration of Helsinki. Written informed consent was obtained from all participants before participation.

### Consent for publication

Consent for publication of de-identified participant data and images was obtained as part of the informed consent process.

### Availability of data and materials

The datasets used and/or analyzed during the current study are available from the corresponding author on reasonable request.

### Authors’ contributions

LW contributed to the study design, performed the data analysis and interpretation, and drafted the manuscript. AMW contributed to the study concept and data acquisition. YZ provided biostatistical guidance. YT and JWA provided scientific guidance and mentorship and contributed to the interpretation of the results and revision of the manuscript. QW contributed to the study design, supervised the research, contributed to the data analysis and interpretation, and critically revised the manuscript. All authors read and approved the final manuscript.

## Acknowledgement

The authors thank Jianing Zhang, Haoze Zhu and William Han for the discussion and advice on the research.

